# Prokaryotic genome expansion is facilitated by phages and plasmids but impaired by CRISPR

**DOI:** 10.1101/474767

**Authors:** Na L. Gao, Jingchao Chen, Martin J Lercher, Wei-Hua Chen

## Abstract

Bacteriophages and plasmids can introduce novel DNA into bacterial cells, thereby creating an opportunity for genome expansion; conversely, CRISPR, the prokaryotic adaptive immune system, which targets and eliminates foreign DNAs, may impair genome expansions. Recent studies presented conflicting results over the impact of CRISPR on genome expansion. In this study, we assembled a comprehensive dataset of prokaryotic genomes and identified their associations with phages and plasmids. We found that genomes associated with phages and/or plasmids were significantly larger than those without, indicating that both phages and plasmids contribute to genome expansion. Genomes were increasingly larger with increasing numbers of associated phages or plasmids. Conversely, genomes with CRISPR systems were significantly smaller than those without, indicating that CRISPR has a negative impact on genome size. These results confirmed that on evolutionary timescales, bacteriophages and plasmids facilitate genome expansion, while CRISPR impairs such a process in prokaryotes. Furthermore, our results also revealed that CRISPR systems show a strong preference for targeting phages over plasmids.

## Introduction

Gene duplication and/or horizontal gene transfer (HGT) play important roles in functional innovation and species adaptation, and are the main sources of genome expansions (1–5). In prokaryotes, it has been shown that the importance of HGT for genome expansions can even outweigh that of gene duplication (6, 7).

Mobile DNA elements such as bacteriophages (referred to as phages below) and plasmids can infect their hosts and introduce novel DNAs into the host genomes (8–11). They often have a very narrow range of hosts; but under certain conditions, such as antibiotic stress, phages and plasmids can expand their host ranges (12). Therefore, phages and plasmids are important sources of HGT and of prokaryotic innovations, and consequently drive bacterial evolution and adaptation (13–15).

Phages and plasmids are widely distributed in prokaryotes. Unlike plasmids, phages are pathogens that often lead to lysis of their hosts (16, 17). Over the course of prokaryotic evolution, bacteria and archaea developed various defense systems against phages, plasmids, and other invading genetic elements (18). CRISPR (clustered regularly interspaced short palindromic repeats), the adaptive immune system of prokaryotes, is a recently recognized player in the ongoing arms race between viruses and hosts, and plays an important role in the dynamic process by which the genomes of prokaryotes and mobile elements coevolve. CRISPR systems are wide-spread in prokaryotes, found in about 40% of bacteria and 90% of archaea (19–22). CRISPR systems can also target plasmids (23), although plasmids are not necessarily detrimental to their host’s fitness but instead often carry a diverse range of antimicrobial and biocide resistance genes that may help their hosts to survive under certain conditions (24, 25).

Based on the above observations, it is reasonable to speculate that over the course of evolution, phages and plasmids may contribute to the expansion of prokaryotic genomes, while CRISPR systems may impair such a process. These speculations are consistent with recent observations that CRISPR limits HGT by targeting foreign DNAs (23, 26). However, controversial observations have also been reported recently. For example, Gophna and colleagues did not observe the expected negative correlation between CRISPR activity in microbes with three independent measures of recent HGT, leading them to conclude that the inhibitory effect of CRISPR against HGT is undetectable (27). Furthermore, a recent study revealed that CRISPR-mediated phage resistance can even enhance HGT by increasing the resistance of transductants against subsequent phage infections (28). These observations appear surprising, as the restricted acquisition of foreign genetic material is believed to be one of the sources of the maintenance fitness cost of CRISPR systems and may be one of the reasons for the patchy distribution of CRISPR among bacteria (29, 30). Thus, it is currently unclear what long-term effects CRISPR, phages, and plasmids have on genome expansion.

In this study, we first collected a comprehensive dataset of prokaryotes and their associations with phages, plasmids, and CRISPR systems. We then applied a generalized linear model to evaluate the contributions of phages, plasmids, and CRISPR to genome size. After controlling for genome GC (guanine+cytosine) content, which is known to correlate significantly with genome size (31, 32), we found that both phages and plasmids are associated with larger genomes, while the presence of a CRISPR system is associated with small genome size. Genome sizes increase with increasing numbers of associated phages and plasmids. Our results clearly indicate that in the long run, phages and plasmids facilitate genome expansions, while CRISPR impairs such a process in prokaryotes. Furthermore, our results also reveal a striking preference of CRISPR systems for targeting phages rather than plasmids, consistent with the typical consequences of phage and plasmid infections to the hosts and the roles of CRISPR as a defense system.

## Results and discussion

### Prokaryotic genomes and their associations with phages, plasmids and CRISPRs

To systematically investigate the impacts of phages, plasmids, and CRISPRs on genome expansion, we assembled a list of 5,994 completely sequenced prokaryotic genomes and obtained their associations with phages, plasmids, and CRISPRs; for details please consult the Materials and Methods section and Supplementary Table 1.

As shown in Figure 1A, we found that 53.98% of prokaryotes had no known associations with infecting phages. 14.88%, 16.68%, and 14.46% of prokaryotes were associated with one, two to three, and more than three phages, respectively (Figure 1A). In addition, we found that 67.46% of prokaryotes did not associate with plasmids, while 14.75%, 11.68%, and 6.12% of the genomes associated with one, two to three, and more than three plasmids, respectively (Figure 1B). A previous study suggested that the genomic GC-contents of phages and plasmids often closely resembles that of their hosts (33, 34); consistent with these previous observations, we obtained correlation coefficient values of 0.972 and 0.970 between the GC-contents of the host genomes and their associated phages and plasmids, respectively (Supplementary Figure 1), confirming the high quality of our association data. We found that in total 42.58% of genomes collected in this study contained either phages or plasmids but not both, while 17.98% of genomes contained both phages and plasmids.

**Figure 1.**
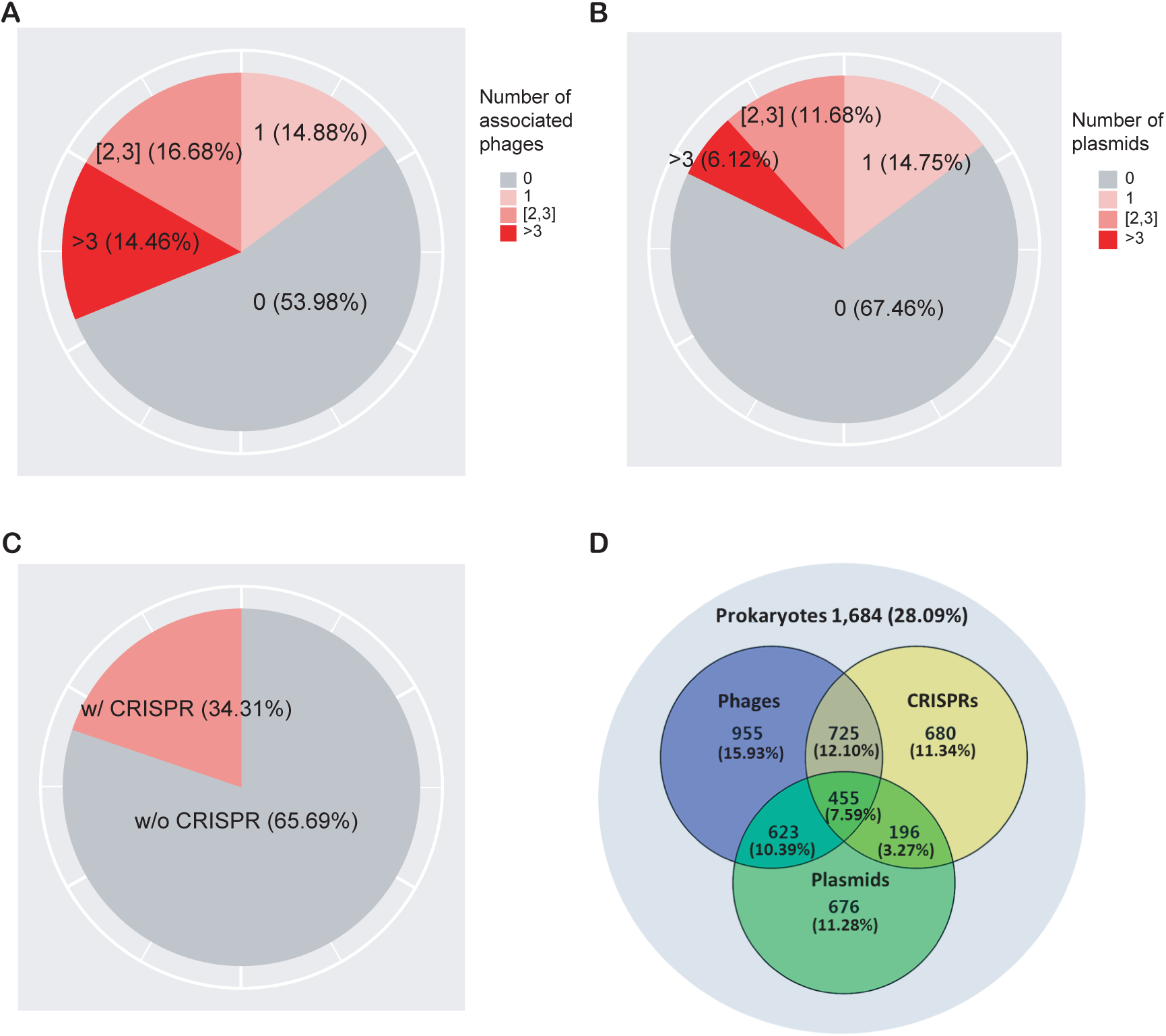
5,994 prokaryotic genomes and their associations with phages (A), plasmids (B), and CRISPRs (C). The Venn diagram (D) shows the overlap of their distributions in prokaryotes. 1,684 genomes (28.09%) were not found to be associated with phages, plasmids, or CRISPRs; 455 (7.59%) genomes were associated with all three elements.

As shown in Figure 1C, we identified CRISPR systems in 34.31% of the prokaryotic genomes (Figure 1C); this percentage is slightly less than previously reported, mainly due to the fact that we removed plasmid-encoded CRISPR systems from our calculation (see Materials and Methods for details). We found that CRISPRs were significantly enriched in phage-associated compared to non-phage-associated genomes (odds ratio OR=1.43, *P*=1.7x10^-14^ from Fisher’s exact test) but not in plasmid-associated compared to non-plasmid-associated genomes (OR=0.96, *P*=0.47). In addition, we found that CRISPRs were more enriched in phage-associated compared to plasmid-associated genomes (OR=2.62, *P*= 9.0x10^-26^, excluding genomes containing both phages and plasmids), suggesting a strong target preferences of CRISPRs toward phages (Table 1).

**Table 1.** Estimated enrichment of CRISPR in phage-associated and plasmid-associated genomes compared to other genomes, and enrichment of CRISPR in phage-associated compared to plasmid-associated genomes.

| Comparison | Odds ratio<sup>b</sup> | *P*-value<sup>c</sup> |
| --- | --- | --- |
| Phage-associated vs. others | 1.43 | 1.75x10<sup>-14</sup> |
| Plasmid-associated vs. others | 0.96 | 0.47 |
| Phage- vs. plasmid-associated | 1.49 | 6.62x10<sup>-11</sup> |
| Phage-associated vs. others<sup>a</sup> | 1.45 | 3.77x10<sup>-11</sup> |
| Plasmid-associated vs. others<sup>a</sup> | 0.56 | 1.01x10<sup>-12</sup> |
| Phage- vs. Plasmid-associated<sup>a</sup> | 2.62 | 8.97x10<sup>-26</sup> |

<sup>a</sup> excluding genomes contained both phages and plasmids.
<sup>b</sup> odds ratio OR > 1 indicates enrichment of CRISPR in the first group, while OR < 1 indicates depletion.
<sup>c</sup> *P*-values indicate whether CRISPR is significantly enriched or depleted in the first group as compared with the second according to Fisher's exact test.

### Phages and plasmids are associated with larger genomes, while CRISPR is associated with smaller genomes

We next investigated which factors contribute significantly to genome size. Previous results have shown a strong correlation between genomic GC content and genome size (31); GC content may even play a causal role in shaping genome size (32). Applying a linear model (LM, see Materials and Methods for details), we found that GC content was indeed the strongest predictor of genome size (Table 2). The LM analysis also revealed that the presence/absence of phages, plasmids, and CRISPR all significantly influenced genome size; the presences of phages and of plasmids were associated with increased genome sizes, while CRISPR was associated with decreased genome sizes (Table 2). We estimated that the relative importance of these factors for genome size were 89% for GC-content, 5.8% for phage presence, 4.4% for plasmid presence, and 0.38% for CRISPR presence. Interestingly, we found that the presence of both phages and plasmids in the same genome was associated with a smaller genome size than expected if the contribution of phages and plastids was independent plasmids (*i.e.*, the interaction term phages*plasmids was negative, Table 2). Unless stated otherwise, we thus limit our further analyses to prokaryotes that contained either phages or plasmids but not both. Note that our conclusions on the influence of phages, plasmids, and CRISPR systems on genome size remain unchanged if we perform separate analyses on genomes containing no phages and on genomes containing no plasmids (Table 2).

**Table 2.** Relative importance of various factors for genome size in a linear model (LM).

| Dataset | Factor | Coefficient | *P*-value | Relative importance |
| --- | --- | --- | --- | --- |
| All | GC% | 0.089 | < 2x10<sup>-16</sup> | 89.51% |
|  | plasmid | 0.754 | < 2x10<sup>-16</sup> | 5.77% |
|  | phage | 0.598 | < 2x10<sup>-16</sup> | 4.35% |
|  | CRISPR | -0.158 | 2.0x10<sup>-4</sup> | 0.38% |
|  | phage*plasmid | -0.216 | 0.012 | - |
| No plasmids | GC% | 0.09 | < 2x10<sup>-16</sup> | 93.93% |
|  | phage | 0.596 | < 2x10<sup>-16</sup> | 5.65% |
|  | CRISPR | -0.164 | 1.1x10<sup>-3</sup> | 0.41% |
| No phages | GC% | 0.093 | < 2x10<sup>-16</sup> | 93.76% |
|  | plasmid | 0.743 | < 2x10<sup>-16</sup> | 6.01% |
|  | CRISPR | -0.145 | 0.024 | 0.28% |

Note: The equation of “All” dataset used in the linear model (LM) is size ~ GC% + plasmid + phage + CRISPR + phage\*plasmid. Here, size represents the genome size;
GC% represents the genomic GC-content of the host genome; plasmid, phage, and CRISPR represent whether the host genomes are associated with plasmids, phages, and CRISPR, respectively. The “Coefficient” column contains estimated regression coefficients calculated by ordinary least squares. Relative importance was calculated using the ‘relaimpo’ package (49); the equation of “No plasmids” dataset is size ~ GC% + phage + CRISPR; and the equation of “No phages” dataset is size ~ GC% + plasmid + CRISPR.

### Increasing numbers of phages and plasmids are associated with increased genome sizes

We next investigated the impact of the numbers of phages and plasmids on genome size. Phages and plasmids often have very narrow host ranges (35); the number of known associations with phages may indicate the ability of the prokaryotic host to acquire external novel DNA. Consistent with our expectation, we found that genomes associated with more phages had larger overall genomes (Figure 2A). We observed similar results with plasmids (Figure 2B).

**Figure 2.**
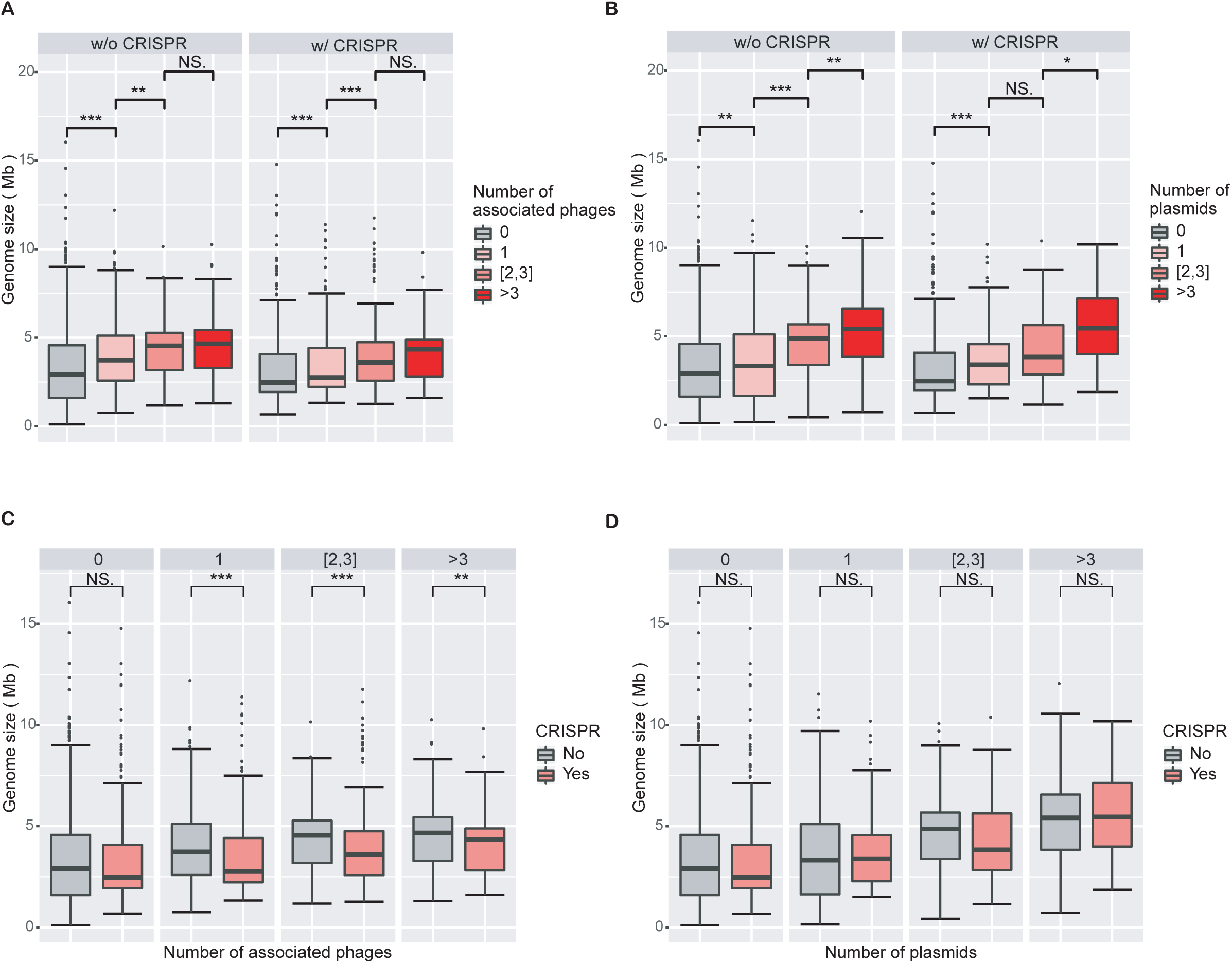
Increasing numbers of phages and plasmids are associated with increased genome sizes, while phage-associated genomes with CRISPR systems are smaller than those without CRISPR systems. A) Boxplot of genomes size as a function of the number of associated phages. Genome sizes are larger with increasing numbers of associated phages, regardless of whether genomes encode CRISPR systems. B) Boxplot of genomes size as a function of the number of associated plasmids. The impact of plasmids on genome size is similar to that of bacteriophages. C) Boxplot of genome size as a function of the presence/absence of CRISPRs in genomes associated with phages. Phage-associated genomes with CRISPR systems are significantly smaller in size than those without CRISPR, regardless of the number of phages they are associated with. D) Boxplots of genome sizes in genomes associated with plasmids as a function of the presence/absence of CRISPRs. CRISPRs have no significant impact on genome sizes in genomes associated with plasmids. Wilcoxon rank sum tests were used to compare between groups. Level of significance: *** *P*<0.001; ** *P*<0.01; * *P*<0.05; NS. *P*≥0.05.

Consistent with the results from the LM analysis, we found that phage-associated genomes are statistically significantly smaller when they encode a CRISPR system compared to when they do not (Figure 2C). However, we did not find a corresponding trend in plasmid-associated genomes (Figure 2D). These results are consistent with the different fitness consequences of phage and plasmid invasions to the prokaryotic hosts. Both phages and plasmids can bring exogenous DNA to prokaryotes and decrease the fitness of their hosts, for example by increasing the burden on the host’s transcription and translation apparatus. However, phages typically cause substantial additional fitness decreases through virion production and assembly and eventually host lysis, while plasmids often carry genes that are beneficial to the survival of their hosts under certain circumstances (36, 37). It is thus likely that the CRISPR systems in prokaryotes are more sensitive to phages than to plasmids. This line of argument is also consistent with our results that the presence of CRISPRs is more enriched in phage-associated than in plasmid-associated genomes.

### The influence of associated phages, plasmids, and CRISPR on genome GC-content

We then investigated which factors contribute significantly to genome GC-content. Consistent with our previous results (LM analysis, Table 2), we found that genome size was indeed the most significant predictor of GC-content, with a relative importance of almost 99% (LM analysis, Table 3). The presence of plasmids also had a significant influence on GC-content, with a relative importance of 1% (Table 3). The presence/absence of phages and CRISPR had no significant influence on GC-content by themselves; surprisingly, however, the presence of phages reduced the influence of plasmid presence on GC content.

**Table 3.** Relative importance of various factors for GC-content (GC%) in a LM.

| Dataset | Factors | Coefficient | P-value | Relative importance |
| --- | --- | --- | --- | --- |
| All | size | 4.12 | $< 2 \times 10^{-16}$ | 98.91% |
| | plasmid | -1.135 | $7.29 \times 10^{-3}$ | 1.03% |
|  | phage | -0.007 | 0.983 | 0.06% |
|  | CRISPR | -0.056 | 0.846 | 0.00% |
|  | phage*plasmid | -1.247 | 0.033 | - |
| No plasmids | size | 4.17 | $< 2 \times 10^{-16}$ | 100% |
|  | phage | -0.074 | 0.829 | 0.00% |
|  | CRISPR | 0.066 | 0.847 | 0.00% |
| No phages | size | 4.16 | $< 2 \times 10^{-16}$ | 99.39% |
|  | plasmid | -1.108 | 0.017 | 0.28% |
|  | CRISPR | 1.108 | 0.011 | 0.32% |

Note: The equation of “All” dataset used in the linear model (LM) is GC% ~ size + plasmid + phage + CRISPR + phage\*plasmid; the equation of “No plasmids” dataset is GC% ~ size + phage + CRISPR; and the equation of “No phages” dataset is GC% ~ size + plasmid + CRISPR.

We also investigated whether these factors contribute significantly to GC-content when genomes contain no phages/plasmids. Unsurprisingly, genome size remained the most significant factor for the prediction of genome GC-content, as shown in Table 3, with a relative importance of around 99%. Analysis of genomes without phage-associations confirmed the small influence of plasmid presence on GC content (Table 3). In addition, analysis of genomes without plasmid-associations revealed a small but statistically significant influence of phage presence on GC-content (Table 3).

As shown in Table 4 and Supplementary Figure 3, we did not find clear and consistent trends in GC-content as a function of the number of associated phages or plasmids. These results indicate that the influence of phages and plasmids on genome GC-content is secondary to the effect of genome size.

**Table 4.** Relative importance of various factors for GC-content (GC%) in a LM.

| Factors | Coefficient | P-value | Relative importance |
| --- | --- | --- | --- |
| size | 4.134 | $< 2 \times 10^{-16}$ | 98.10% |
| plasmidNumber | -0.638 | $4.31 \times 10^{-15}$ | 1.67% |
| phageNumber | -0.037 | $3.44 \times 10^{-3}$ | 0.23% |
| CRISPR | -0.131 | 0.646 | 0.01% |

Note: The equation used in the linear model (LM) is $GC\% \sim size + plasmidNumber + phageNumber + CRISPR$ . Here, *plasmidNumber* and *phageNumber* represent the number of plasmids and phages in genomes, respectively.

## Conclusion

We expected that phages and plasmids could facilitate genome expansions because they can bring novel DNAs into prokaryotic cells that can be integrated into the host genome, while CRISPR immune systems could impair such a process by targeting and eliminating foreign DNAs. However, recent studies presented inconsistent results regarding this topic (22, 23, 26–28).

To address this issue, we assembled a comprehensive dataset of prokaryotic genomes and their associations with phages and plasmids. By dividing genomes into distinct groups according to whether they associated with phages and/or plasmids and/or contained CRISPRs, we revealed that genomes with phages or with plasmids were significantly larger than those without, and genome sizes increased with increasing numbers of associated phages/plasmids. Conversely, phage-associated (but not plasmid-associate) genomes with CRISPRs were significantly smaller in size than those without, regardless of the number of associated phages. These results confirm that in the long run, bacteriophages and plasmids facilitate genome expansions while CRISPR impairs phage-driven genome expansions.

Genome size evolution has previously been reported to be associated with that of genomic GC-content (38). Thus, it appeared possible that phage-and/or plasmid-association has a direct effect not only on genome size but also on GC-content. However, in this study, we found only minor influences of phages (Table 6) and plasmids (Tables 5, 7) on genomic GC-content.

Our results also imply that CRISPR immune systems might be more sensitive towards invading phages than plasmids, consistent with the differential fitness burdens brought by the two types of foreign invaders to the hosts (37, 39–41).

Our results differ significantly from several previous studies (27, 28). For example, Gophna *et al.* reported that the inhibitory effect of CRISPR against HGT is undetectable using three independent measures of recent HGT (27). However, it is known that CRISPR spacers – which were used by Gophna *et al*. to assess CRISPR activity (27) – have very high turnover rates, on the time-scale of days (42–44), while HGT genes may take a very long time to be incorporated into existing gene networks (45), suggesting that it is only possible to look at the impacts of CRISPRs on HGTs at evolutionary scales. Interestingly, Gophna *et al.* also studied spacer acquisition and concluded there was a bias toward frequently encountered invasive exogenous genetic elements, especially infecting viruses (27); this is consistent with our conclusion that CRISPRs tend to be more sensitive towards invading phages than plasmids. Recently, Watson *et al.* reported that the CRISPR system of the bacterium *Pectobacterium atrosepticum* enabled the host to resist phage infection, but that this enhanced rather than impeded HGT by transduction (28). However, it is yet to be seen whether or not this phenomenon is unique to *P. atrosepticum*.

## Materials and Methods

### Data

We obtained data from three sources. Microbe-phage interaction data was collected from the MVP database, which we described in a previous publication (35). MVP is one of the latest and largest databases about microbe-phage interactions, which supplied 26,572 interactions between 9,245 prokaryotes and 18,608 viral clusters based on 30,321 evidence entries (35).

The basic genome information from complete archaeal and bacterial genomes, including the number of associated plasmids, was downloaded from the NCBI Genome database (https://www.ncbi.nlm.nih.gov/genome/; accessed on June 28, 2018) (46). The genome size and GC-content from 10,686 complete prokaryotic genomes (287 archaeal and 10,279 bacterial genomes) were identified. 2,827 prokaryotes were associated with plasmids.

The CRISPRs data was obtained from the CRISPRdb database (47) (http://crispr.i2bc.paris-saclay.fr/; last update May 09, 2017). 202 archaeal and 3,059 bacterial genomes were associated with CRISPR systems. 77 of these encode CRISPR on both plasmids and genome, while only 36 encode CRISPR exclusively on plasmids. The 77 genomes which contained plasmid-encoded CRISPR systems were removed from all analyses.

In total, 5,994 prokaryotes were found in both of the first two datasets; among these, 1,950 contained plasmids, 2,758 contained phages, and 2,056 contained CRISPRs on their chromosomes. Detailed information on the dataset can be found in Supplementary Table 1.

### Statistical analysis

All data were analyzed using R v3.4 (48). All pair-wise comparisons between two groups of numeric data (genome sizes or genomic GC-contents) were performed by Wilcoxon rank-sum tests. Linear model (LM) analysis was performed with the R function glm(). Relative importance analysis was performed with the calc.relimp() function available from the R package ‘relatimpo’ (49).

## References

1 Smith G, Macias-Munoz A, Briscoe AD. 2016. Gene Duplication and Gene Expression Changes Play a Role in the Evolution of Candidate Pollen Feeding Genes in Heliconius Butterflies. Genome biology and evolution 8:2581–2596.

2 Nyvltova E, Stairs CW, Hrdy I, Ridl J, Mach J, Paces J, Roger AJ, Tachezy J. 2015. Lateral gene transfer and gene duplication played a key role in the evolution of Mastigamoeba balamuthi hydrogenosomes. Molecular biology and evolution 32:1039–1055.

3 Tsai YM, Chang A, Kuo CH. 2018. Horizontal Gene Acquisitions Contributed to Genome Expansion in Insect-Symbiotic Spiroplasma clarkii. Genome biology and evolution 10:1526–1532.

4 Isambert H, Stein RR. 2009. On the need for widespread horizontal gene transfers under genome size constraint. Biology direct 4:28.

5 Schonknecht G, Chen WH, Ternes CM, Barbier GG, Shrestha RP, Stanke M, Brautigam A, Baker BJ, Banfield JF, Garavito RM, Carr K, Wilkerson C, Rensing SA, Gagneul D, Dickenson NE, Oesterhelt C, Lercher MJ, Weber AP. 2013. Gene transfer from bacteria and archaea facilitated evolution of an extremophilic eukaryote. Science 339:1207–1210.

6 Pal C, Papp B, Lercher MJ. 2005. Adaptive evolution of bacterial metabolic networks by horizontal gene transfer. Nature genetics 37:1372–1375.

7 Treangen TJ, Rocha EP. 2011. Horizontal transfer, not duplication, drives the expansion of protein families in prokaryotes. PLoS genetics 7:e1001284.

8 Malachowa N, DeLeo FR. 2010. Mobile genetic elements of Staphylococcus aureus. Cellular and molecular life sciences: CMLS 67:3057–3071.

9 Yamaguchi T, Hayashi T, Takami H, Ohnishi M, Murata T, Nakayama K, Asakawa K, Ohara M, Komatsuzawa H, Sugai M. 2001. Complete nucleotide sequence of a Staphylococcus aureus exfoliative toxin B plasmid and identification of a novel ADP-ribosyltransferase, EDIN-C. Infection and immunity 69:7760–7771.

10 Jensen SO, Lyon BR. 2009. Genetics of antimicrobial resistance in Staphylococcus aureus. Future microbiology 4:565–582.

11 Lindsay JA. 2010. Genomic variation and evolution of Staphylococcus aureus. International journal of medical microbiology: IJMM 300:98–103.

12 Modi SR, Lee HH, Spina CS, Collins JJ. 2013. Antibiotic treatment expands the resistance reservoir and ecological network of the phage metagenome. Nature 499:219–222.

13 Koonin EV, Wolf YI. 2008. Genomics of bacteria and archaea: the emerging dynamic view of the prokaryotic world. Nucleic acids research 36:6688–6719.

14 Argov T, Azulay G, Pasechnek A, Stadnyuk O, Ran-Sapir S, Borovok I, Sigal N, Herskovits AA. 2017. Temperate bacteriophages as regulators of host behavior. Current opinion in microbiology 38:81–87.

15 Nogueira T, Rankin DJ, Touchon M, Taddei F, Brown SP, Rocha EP. 2009. Horizontal gene transfer of the secretome drives the evolution of bacterial cooperation and virulence. Current biology: CB 19:1683–1691.

16 Deresinski S. 2009. Bacteriophage therapy: exploiting smaller fleas. Clinical infectious diseases: an official publication of the Infectious Diseases Society of America 48:1096–1101.

17 Wernicki A, Nowaczek A, Urban-Chmiel R. 2017. Bacteriophage therapy to combat bacterial infections in poultry. Virology journal 14:179.

18 Luk AW, Williams TJ, Erdmann S, Papke RT, Cavicchioli R. 2014. Viruses of haloarchaea. Life 4:681–715.

19 Huang Q, Luo H, Liu M, Zeng J, Abdalla AE, Duan X, Li Q, Xie J. 2016. The effect of Mycobacterium tuberculosis CRISPR-associated Cas2 (Rv2816c) on stress response genes expression, morphology and macrophage survival of Mycobacterium smegmatis. Infection, genetics and evolution: journal of molecular epidemiology and evolutionary genetics in infectious diseases 40:295–301.

20 Godde JS, Bickerton A. 2006. The repetitive DNA elements called CRISPRs and their associated genes: evidence of horizontal transfer among prokaryotes. Journal of molecular evolution 62:718–729.

21 Seed KD, Lazinski DW, Calderwood SB, Camilli A. 2013. A bacteriophage encodes its own CRISPR/Cas adaptive response to evade host innate immunity. Nature 494:489–491.

22 Makarova KS, Haft DH, Barrangou R, Brouns SJ, Charpentier E, Horvath P, Moineau S, Mojica FJ, Wolf YI, Yakunin AF, van der Oost J, Koonin EV. 2011. Evolution and classification of the CRISPR-Cas systems. Nature reviews. Microbiology 9:467–477.

23 Marraffini LA, Sontheimer EJ. 2008. CRISPR interference limits horizontal gene transfer in staphylococci by targeting DNA. Science 322:1843–1845.

24 McCarthy AJ, Lindsay JA. 2012. The distribution of plasmids that carry virulence and resistance genes in Staphylococcus aureus is lineage associated. BMC microbiology 12:104.

25 Shabbir MA, Hao H, Shabbir MZ, Wu Q, Sattar A, Yuan Z. 2016. Bacteria vs. Bacteriophages: Parallel Evolution of Immune Arsenals. Frontiers in microbiology 7:1292.

26 Bikard D, Hatoum-Aslan A, Mucida D, Marraffini LA. 2012. CRISPR interference can prevent natural transformation and virulence acquisition during in vivo bacterial infection. Cell host & microbe 12:177–186.

27 Gophna U, Kristensen DM, Wolf YI, Popa O, Drevet C, Koonin EV. 2015. No evidence of inhibition of horizontal gene transfer by CRISPR-Cas on evolutionary timescales. The ISME journal 9:2021–2027.

28 Watson BNJ, Staals RHJ, Fineran PC. 2018. CRISPR-Cas-Mediated Phage Resistance Enhances Horizontal Gene Transfer by Transduction. mBio 9.

29 Baltrus DA. 2013. Exploring the costs of horizontal gene transfer. Trends in ecology & evolution 28:489–495.

30 Frost LS, Leplae R, Summers AO, Toussaint A. 2005. Mobile genetic elements: the agents of open source evolution. Nature reviews. Microbiology 3:722–732.

31 Chen W-H, van Noort V, Lluch-Senar M, Hennrich ML, H. Wodke JA, Yus E, Alibés A, Roma G, Mende DR, Pesavento C, Typas A, Gavin A-C, Serrano L, Bork P. 2016. Integration of multi-omics data of a genome-reduced bacterium: Prevalence of post-transcriptional regulation and its correlation with protein abundances. Nucleic Acids Research 44:1192–1202.

32 Chen WH, Lu G, Bork P, Hu S, Lercher MJ. 2016. Energy efficiency trade-offs drive nucleotide usage in transcribed regions. Nature communications 7:11334.

33 Hiroshi Nakashima KHaKM. 2015. Relationship of Genomic G+C Content between Phages/Plasmids and Their Hosts. British Biotechnology Journal:9(1): 1–9.

34 Ren J, Ahlgren NA, Lu YY, Fuhrman JA, Sun F. 2017. VirFinder: a novel k-mer based tool for identifying viral sequences from assembled metagenomic data. Microbiome 5:69.

35 Gao NL, Zhang C, Zhang Z, Hu S, Lercher MJ, Zhao X-M, Bork P, Liu Z, Chen W-H. 2018. MVP: a microbe-phage interaction database. Nucleic Acids Research 46:D700–D707.

36 Dionisio F, Conceicao IC, Marques AC, Fernandes L, Gordo I. 2005. The evolution of a conjugative plasmid and its ability to increase bacterial fitness. Biology letters 1:250–252.

37 Jiang W, Maniv I, Arain F, Wang Y, Levin BR, Marraffini LA. 2013. Dealing with the evolutionary downside of CRISPR immunity: bacteria and beneficial plasmids. PLoS genetics 9:e1003844.

38 Gao N, Lu G, Lercher MJ, Chen WH. 2017. Selection for energy efficiency drives strand-biased gene distribution in prokaryotes. Scientific reports 7:10572.

39 Weinberger AD, Sun CL, Plucinski MM, Denef VJ, Thomas BC, Horvath P, Barrangou R, Gilmore MS, Getz WM, Banfield JF. 2012. Persisting viral sequences shape microbial CRISPR-based immunity. PLoS computational biology 8:e1002475.

40 Canchaya C, Fournous G, Brussow H. 2004. The impact of prophages on bacterial chromosomes. Molecular microbiology 53:9–18.

41 Pleska M, Guet CC. 2017. Effects of mutations in phage restriction sites during escape from restriction-modification. Biology letters 13.

42 Deveau H, Barrangou R, Garneau JE, Labonte J, Fremaux C, Boyaval P, Romero DA, Horvath P, Moineau S. 2008. Phage response to CRISPR-encoded resistance in Streptococcus thermophilus. Journal of bacteriology 190:1390–1400.

43 Horvath P, Romero DA, Coute-Monvoisin AC, Richards M, Deveau H, Moineau S, Boyaval P, Fremaux C, Barrangou R. 2008. Diversity, activity, and evolution of CRISPR loci in Streptococcus thermophilus. Journal of bacteriology 190:1401–1412.

44 Tyson GW, Banfield JF. 2008. Rapidly evolving CRISPRs implicated in acquired resistance of microorganisms to viruses. Environmental microbiology 10:200–207.

45 Lercher MJ, Pal C. 2008. Integration of horizontally transferred genes into regulatory interaction networks takes many million years. Mol Biol Evol 25:559–567.

46 NCBI Resource Coordinators. 2018. Database resources of the National Center for Biotechnology Information. Nucleic acids research 46:D8–D13.

47 Grissa I, Vergnaud G, Pourcel C. 2007. The CRISPRdb database and tools to display CRISPRs and to generate dictionaries of spacers and repeats. BMC bioinformatics 8:172.

48 R core Team. 2017. R: A Language and Environment for Statistical Computing.

49 U G. 2006. Relative importance for linear regression in R: The package relaimpo. J. Stat. Softw:17:11–27.

